# Gender identity and sexual experience affect mating behaviour and chemical profile in the lesser mealworm, *Alphitobius diaperinus* (Coleoptera: Tenebrionidae)

**DOI:** 10.1101/2021.08.04.455096

**Authors:** Erika Calla-Quispe, Carlos Martel, Alfredo J. Ibañez

**Affiliations:** Instituto de Ciencias Ómicas y Biotecnología Aplicada, Pontificia Universidad Católica del Perú, Av. Universitaria 1801, San Miguel 15088, Lima, Peru

## Abstract

*Alphitobius diaperinus* (Coleoptera: Tenebrionidae), the lesser mealworm, is one of the most significant pests of the poultry industry worldwide. These insects cause structural damage in poultry houses and transmit several diseases, impacting chickens’ productivity and rearing costs. Although semiochemicals may offer alternative insect pest management strategies, basic information regarding pheromone identity and their role on the behavioural ecology according to their circadian pattern of sexual behaviour of *A. diaperinus* is essentially lacking. This study is aimed to analyse the relation of gender identity and sexual experience of adults of *A. diaperinus* on their mating behaviour and whether this response is related to their CHC profiles secreted. The following steps were taken to achieve the study’s goal. First, the circadian pattern of their sexual activity was observed in newly emerged pairs for at least twenty-one days (virgin adults) and experienced adults collected from the field to identify a difference based on their sexual experience and achieve the optimal mating season to develop the following assays. Subsequently, Y-tube olfactometer bioassays were conducted to evaluate their odour bouquet attraction based on gender and sexual experience. Additionally, mating behaviour bioassays were conducted to evaluate the two factor effects. Finally, cuticular analysis was performed using gas chromatography-mass spectrometry to evaluate possible chemical differences based on the two factors. With statistical and multivariate analysis, we found that behavioural, mating and chemical responses are different based on their sexual experience. The mating sequences were described into precopulatory, copulatory and postcopulatory phases. This finding gives us a deeper understanding of the sexual communication during mating. In summary, our findings provide new insights into the mating system and chemical ecology of *A. diaperinus*. The results presented here may serve as a base for further studies to develop strategies for managing this pest.

## Introduction

*Alphitobius diaperinus* (Panzer, 1797) (Coleoptera: Tenebrionidae), commonly known as the lesser mealworm, has long captured the attention of the people involved in the poultry industry due to being a worldwide pest in poultry farms (1–3). Individuals of this species grow in the chicken litter, within a mixture of wasted feed, faeces and feathers. They can structurally damage the poultry houses (1,4–7), which generates economic losses. They also act as vectors of several viral (8,9), fungal (10–12) and bacterial (13–16) diseases, which can cause poultry weight loss and even death (3,17–19). To control this pest, poultry farmers invest in pesticides and drugs, i.e., to prevent diseases on chickens (20,21). Nevertheless, the widespread use of pesticides (e.g., carbamates, organophosphates and pyrethroids) may cause resistant insect populations, harm birds, and lead to poultry house contamination (1,2,4,22–28). Consequently, alternative methods are researched to reduce the use of insecticides. These alternative methods may include physical methods (23,29,30), biological control (11,22,31–35) and chemical methods using natural products (18,36) such as bioinsecticides (28) or semiochemicals (5,17,24,37–41).

Semiochemicals are particularly effective as biological control, environmentally friendly, and highly specific based on pheromone composition (37,42–44). Recently, some studies have addressed the identification of aggregation and alarm pheromones from *A*. *diaperinus* (5,38,39,41). Nevertheless, the identification of the pheromones alone does not warrant the success in controlling the *A. diaperinus* population since insect responses are heavily context-dependent (e.g., environmental factors (45–47), interspecific interaction (48,49), population origin (17,24,40,50). Furthermore, as most organisms followed physiological and behavioural changes to a 24-hour cycle (i.e., circadian pattern), circadian patterns of insect behaviour and particularly reproductive activity can affect the performance of semiochemicals and pheromones (51–56). Therefore, circadian behaviour is a crucial component of almost any ecological and evolutionary process (55,57). Thus, basic knowledge such as identifying processes affecting aggregation or mating behaviour addressing their circadian pattern of sexual behaviour is crucial for more efficient pest management (44,57–62).

To achieve sexual reproduction, insect adults develop a set of behavioural displays (i.e., mating behaviour), which involves partner recognition, courtship, and copulation (44,63,64). In this regard, sexual communication in insects is based on visual, olfactory, tactile and even auditory stimuli (62,65,66). From those mentioned, olfactory stimuli based on chemicals are by far the most critical signals (64,67–69). Chemical signalling in sexual behaviour systems involves highly volatile compounds that promote attractiveness over long distances and compounds that can influence close-range orientation; these chemicals are perceived by direct antennal contact with the insect cuticles (58,60,67,68). Insect cuticles are made by a lipid wax layer, in which cuticular hydrocarbons (CHC) are a significant contributor (49,70,71). Besides their function of limiting water loss and reducing external damage by toxins and pathogens, CHCs also play a critical role in chemical communication for species, nest or mate recognition, and signalling reproductive status (49,61,62,64,70,72–74). In beetles, volatiles associated with intra-specific long attraction is usually associated with aggregation pheromones; in contrast, low volatile and non-volatile compounds are associated with contact sex pheromones (58,72,75,76). CHC compounds are shared by males and female adults in some beetle species (61,70,76), which explain the observed homosexual behaviour upon antennal contact, e.g., in some Scarabaeidae, Silphidae and Tenebrionidae beetles (68,70,77–79).

Since homosexual behaviour is known in *A. diaperinus* (38,40,41), it is expected that both genders share part of their CHC profiles. It has been shown that aggregation pheromones are produced by virgin males of *A. diaperinus*, which attracts males and females (5). However, it is unknown whether this attraction can be affected by gender identity or sexual experience. Furthermore, we expect gender identity and sexual experience in a circadian context can impact semiochemical and pheromone activity. Given that insect CHC profiles are associated with gender identity and sexual experience (61,70,73,80,81), we were interested in finding out whether or not these two factors affect mating behaviour responses and, if so, could those differential behaviours be related to differences associated with their chemical profiles? To answer these questions, we first identified the circadian sexual behaviour of *A. diaperinus* adults. We then tested the attractiveness and behavioural mating responses of males and females with different sexual experiences to different CHC profiles of other *A. diaperinus* adults.

## Materials and methods

### Studied species

Adults of *A. diaperinus* were obtained from a commercial poultry production located in the surroundings of Lima (12°09′27.8″ S 76°53′46.2″ W). All four live stages of *A. diaperinus* were reared in our laboratory in aquarium glass boxes (30 × 25 × 20 cm). In one of our glass boxes, hundreds of male and female adults were kept and let free to reproduce; new emerging larvae were then removed and transferred to a second glass box. To facilitate the handling of the larvae, they were kept in groups within Petri dishes. This second box was monitored twice a week to feed them and to find pupae. Each new pupa was sexed (following (82); i.e., females “F” and males “M”), and then isolated in a Petri dish to avoid they mate in order to obtain non-mated adults. In case of gender recognition could not be carried out at pupa stage (e.g., first collected adults), gender recognition was carried out by pressing their protracted ventral abdomen to see the genitalia under a stereomicroscope. Larvae and adults of *A. diaperinus* were fed using commercial wheat flour “Blanca flor” (Alicorp, Lima, Peru) and tap water, which was provided by using a humidifying towel paper within the boxes. Glass boxes were kept in an environmentally controlled climate chamber (Memmert HPP750, Memmert GmbH, Schwabach, Germany) at a constant temperature of 30 °C and humidity of 50 %, with a photoperiod of 12:12 h (light : dark) (photophase 07:00 to 19:00 h and scotophase 19:00 to 07:00 h; GMT-5). Adults being 21-d-old or older were used to guarantee sexual maturity. Non-mated adults were considered as sexually inexperienced (referred to as ‘virgin’ [v]), whereas adults exposed to high densities of males and females were catalogued as experienced (‘exp’). Fig 1 shows an overall workflow of the different steps of this study, which are described below. No insects were harm during experiments and when killed, it was instantly by freezing.

**Fig 1.**
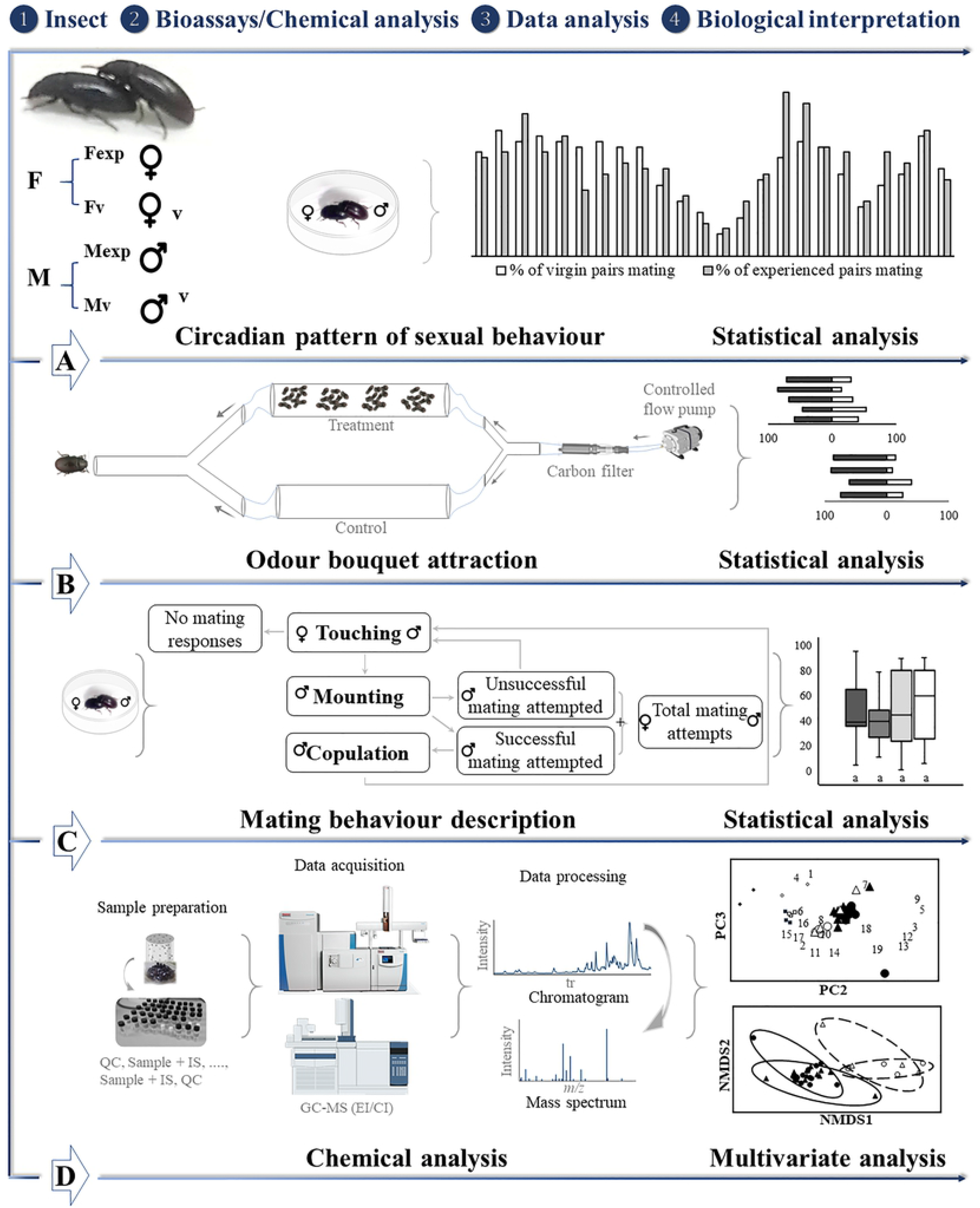
An integrative behavioural and chemical perspective for the analysis of mating behaviour of adults of *Alphitobius diaperinus* based on their gender identity and sexual experience. (A) Circadian pattern of sexual activity, (B) CHC attractiveness, (C) mating behaviour and (D) chemical profile analysis based on CHC affected by gender and sexual experience.

### Circadian pattern of sexual activity

We recorded the mating behaviour of adult pairs (i.e., one female and one male) for 15 min each hour along the day for 5 days. The observations were performed over a Petri dish (60 × 15 mm), and a total of 10 pairs were utilized each day, totalizing 50 adult pairs. When a male exposed his genitalia while mounting a female, it was classified as a matting attempt. Males were not allowed to accomplish mating to continue observations along the day. The proportion of mating attempts was then calculated at each hour. During periods of darkness, observations were performed under an incandescent bulb with red light. One hour prior and between observations, experienced adults were individually kept in Petri dishes to increase sexual desire. After each observation, Petri dishes were washed with methanol.

### CHC attractiveness based on gender and sexual experience

A three-arm olfactometer (i.e., Y-tube, 8 mm thickness, 10 cm apparatus trunk followed by 5 cm length arm) was used to test the attractiveness of CHC compounds of males and females. Each Y-tube arm was connected to one of the five different treatments, which consisted of a glass chamber bearing one of the following options: (i) an empty control (C), (ii) 20 experienced females (Fexp), (iii) 20 virgin females (Fv), (iv) 20 experienced males (Mexp), or (v) 20 virgin males (Mv) (see Fig 1a). The odour released by each treatment was then driven to the olfactometer by an air compressor that provided an airflow at a constant rate of 0.3 L/min through activated charcoal filters. A virgin male or female was placed within the trunk of the olfactometer. Then, it was observed which arm the insect chose within a time of up to 5 min. The beetle odour decision was registered when the beetle entered at least 1 cm into one of the arms. All tested beetles were used only once, whereas beetles within the chambers were allowed to acclimatize for 1 h before the tests to avoid them producing unattractive compounds such as alarm pheromones. The assignment of odour sources to each arm was reversed after each trial to avoid potential illumination and directional bias (59). After each test, the olfactometer and chambers were washed with methanol. All trials were carried out at the peak of circadian sexual behaviour (S1 Fig).

### Effect of gender and sexual experience on mating behaviour

Behavioural experiments with treatments based on all combinations of gender and sexual experience were prepared (i.e., Fv–Mv, Fv–Mexp, Fexp–Mv, Fexp–Mexp). Each pair (i.e., one male and one female) of adults of *A. diaperinus* was placed in a Petri dish (60 × 15 mm) containing a filter paper at the inner bottom and observed for a total of 10 min. The following behaviours developed during mating attempts were timed: touching (i.e., time in touching the other insect’s cuticle with their antennae or their prothoracic leg), mounting (i.e., time upon a male is over and bends his abdomen exposing the genital organ until mating), and copulation (i.e., time in copula). The recorded time per response per trial was then standardised by using a percentage scale (i.e., total trial time was transformed to 100 %). Additionally, the number of times of successful mating attempts (i.e., copulation occurred), unsuccessful mating attempts (i.e., unsuccessful copulation), and total mating attempts (i.e., total successful and unsuccessful mating attempts) were recorded. The recorded number of times per trial was also standardised using a percentage scale based on the total mating attempts.

### Chemical analyses of CHC profiles

Twelve individuals from each group based on gender and sexual experience (i.e., Fexp, Fv, Mexp, Mv) were killed by freezing, thawed for 10 min at room temperature. Their CHCs were extracted by immersing them in 2 mL hexane (MS grade, Sigma-Aldrich) and shaking in a vortex for 2 min at 2000 rpm. For quantitative analysis, 1 μL of 1000 ppm hexadecane (C16) was added as an internal standard in all the extracts. Later, CHC extracts were concentrated to 120 μL using a gentle stream of nitrogen and then stored at −20 °C for subsequent chemical analysis.

Extracts were analysed by gas chromatography (GC) coupled to quadrupole time-of-flight mass spectrometry (MS; Agilent 7250 GC/Q-TOF, Santa Clara/CA, USA) with electron and chemical ionization (EI and CI, respectively). GC was equipped with a DB-5 column (30 m × 0.25 mm i.d., 0.25 μm thickness film). Aliquots of 1 μL cuticular extract were injected in the GC, which was programmed at an oven temperature of 50 °C for 2 min, increased at 10 °C/min until 250 °C, and hold for 20 min. Injections were made in splitless mode with helium as the carrier gas (1.5 mL/min), injector temperature at 250 °C, and detector temperature at 270 °C. Subsequently, the same extracts were measured using a GC-MS with EI at 70 eV with a scan range from *m/z* 50–750. Data mining was done using MS-DIAL v4.6., which provided peak alignments of data based on total ion current of GC-MS with CI analyses (83). After that, filtering was performed using principal component analysis (i.e., PCA of log10-transformed and Z-score-normalized data) and one-way ANOVA (*p*-anova<0.01) through MATLAB vR2019b. This filtering allowed us to track data quality, reduce the data dimensionality, identify potential outliers in the dataset, and identify sample clusters (84–87). To final filtering of each feature was considered the following threshold: that the average area of the sample was at least three times the average area of the C16 and the blank. After data curation, to ensure the identification of compounds, selected samples were analysed by gas chromatography coupled to an APPI-Q-Exactive HF mass spectrometer (Thermo Fisher Scientific, USA) following the methodology explained above. Identification of CHC compounds was achieved by comparing mass spectra and retention indices of unidentified compounds with commercial databases such as PubChem, Fiehn BinBase, MoNA volatile, Chemspider, Metlin and NIST.

### Statistical analysis

Differences in mating activity throughout a circadian cycle and differences in the total time of male mating reactions (i.e., touching, mounting, copulation and mating time), as well as the total mating attempts (i.e., successful, unsuccessful and total mating attempts), among all tested treatments, were performed by non-parametric Kruskal-Wallis tests, followed of pairwise comparisons by Mann-Whitney-U tests with a Bonferroni correction. Choices made by *A*. *diaperinus* adults in the olfactometer bioassays were analysed by exact binomial tests (50 % chance of selecting each arm). All these statistical tests were performed using the *stats* package of the R software (R version 3.6.1 (88)).

A non-metric multidimensional scaling (NMDS) was performed to display the dissimilarities among groups graphically. NMDS ordination was based on Bray–Curtis similarity of the square-root transformed dataset. Furthermore, to identify statistical differences in the CHC profiles among groups, a permutational multivariate analysis of variance (PERMANOVA) was run based on Bray-Curtis similarity of a square-root transformed dataset. A total of 99999 permutations and a Holm correction was carried out, considering “gender” as a fixed factor and “sexual experience” as a nested factor within “gender”. The CHC dataset was based on the relative proportions of all the identified compounds (see results). NMDS and PERMANOVA were performed using the *vegan* package (89), whereas Holm correction using the *RVAideMemoire* package (90) of R.

## Results

### Circadian pattern of sexual activity

Independently of their sexual experience (i.e., virgin or experienced), the mating behaviour of *A. diaperinus* adults was displayed throughout the day (S1 Fig). On the one hand, virgin adults showed more mating activity between 2 h and 9 h after the light was on (i.e., photophase) and at 17 h to 18 h and 23 h, under dark conditions (i.e., scotophase). In contrast, the lowest mating activity was recorded between 13 h and 14 h (S1 Fig). On the other hand, experienced adults were more sexually active between the 3 h and 5 h on the photophase (S1 Fig), and at 8 h to 11 h on the scotophase (S1 Fig); in contrast, the lowest activity was also observed between 13 h and 14 h (S1 Fig). Nevertheless, mating behaviour activity was only significantly higher between 16 h and 17 h for both virgin and experienced adults (S1 Fig, Mann-Whitney-U test with Bonferroni correction, p<0.05).

### Attractiveness based on gender and sexual experience

Fv were significantly more attracted to CHC odour of Mv (Fig 2; Binomial test, p<0.05, n = 31), Mexp (Fig 2; Binomial test, p<0.01, n = 34) and, barely significantly, of Fv (Fig 2; Binomial test, p<0.07, n = 31) over control treatment. However, Fv did not show any preference when tested to CHC odour of Fexp against control or Fexp against Mexp (Fig 2; Binomial test, both p>0.05). In addition, Mv were significantly more attracted to CHC odour of Mv (Fig 2; Binomial test, p<0.001, n = 30), Fexp (Fig 2; Binomial test, p<0.001, n = 40) and, barely significantly, of Fv (Fig 2; Binomial test, p<0.1, n = 53) compared to control. However, due to not enough Mv individuals, no experiments were carried out.

**Fig 2.**
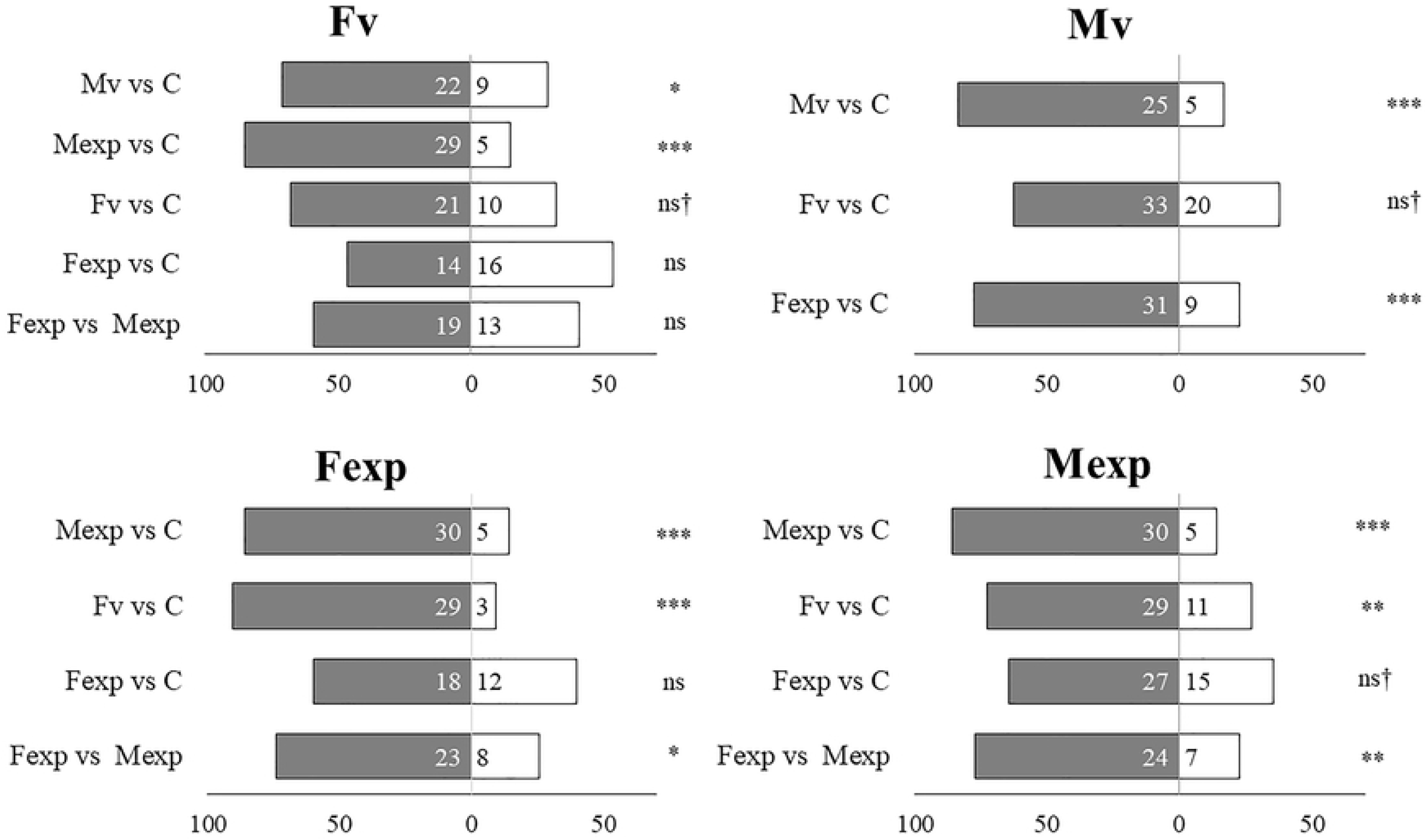
The attractiveness of *Alphitobius diaperinus* adults based on CHC odour by gender and sexual experience. Significant difference (* p<0.05; ** p<0.01; *** p<0.001) and ns=non-significant according to the exact binomial test.

In the case of Fexp, they were significantly more attracted to Mexp (Fig 2; Binomial test, p<0.001, n = 35) and Fv (Fig 2; Binomial test, p<0.001, n = 32) odour over control, as well as, significantly attracted to Fexp scent over Mexp (Fig 2; Binomial test, p<0.05, n = 31). However, they did not show any preference when tested Fexp against control (Fig 2; Binomial test, p>0.05, n = 30). Additionally, Mexp were significantly more attracted to Mexp (Fig 2; Binomial test, p<0.001, n = 35) and Fv (Fig 2; Binomial test, p<0.01, n = 40), as well as significantly attracted to Fexp odour over Mexp (Fig 2; Binomial test, p<0.01, n = 31) and barely significantly attracted to Fexp compared to control (Fig 2; Binomial test, p<0.09, n = 32).

### CHC attractiveness based on gender and sexual experience

The three distinct phases of mating behaviour in *A. diaperinus* are summarized in Fig 1c. The precopulatory phase (i.e., during the insect localization) began when both adults approached between them. After approaching, they touched their cuticles with their antennae or their prothoracic leg. When the male recognised the female, he attempts to mount her from the back, bending his abdomen and exposing the genital organ. Females also contributed to both adults have a successful attempt mating while placing herself under the male and exposing her genitalia by aperture the last sternite. In the copulatory phase, the male grasped the female’s cuticle with his prothoracic and mesothoracic leg and touched her prothorax with his antenna and maxillary palp. Interestingly, copula was on average accomplished only once per trial. In the postcopulatory phase, both adults kept touching and stay that way. When a male did not have a successful mating due to his small size related to the female, both insects attempted mating again (S1 Table).

The Kruskal–Wallis test shows that the touching responses of *A. diaperinus* adults were not significantly different among treatments (Fig 3, Table 1; p>0.05). On the contrary, the pattern of mounting and copulation responses was quite different among treatments (Table 1; Kruskal-Wallis test; p<0.0001). Specifically, Fv-Mv and Fexp-Mv resulted in significantly longer mounting than Fv-Mexp (Fig 3; Mann-Whitney-U test with Bonferroni correction, p<0.05); whereas we did not find differences in mounting in Fexp-Mexp compared to any other treatment. However, we did find significant differences in copulation when comparing Fv-Mv and Fexp-Mv against Fv-Mexp and Fexp-Mexp (Fig 3; Mann-Whitney-U test with Bonferroni correction, p<0.05 for all).

**Fig 3.**
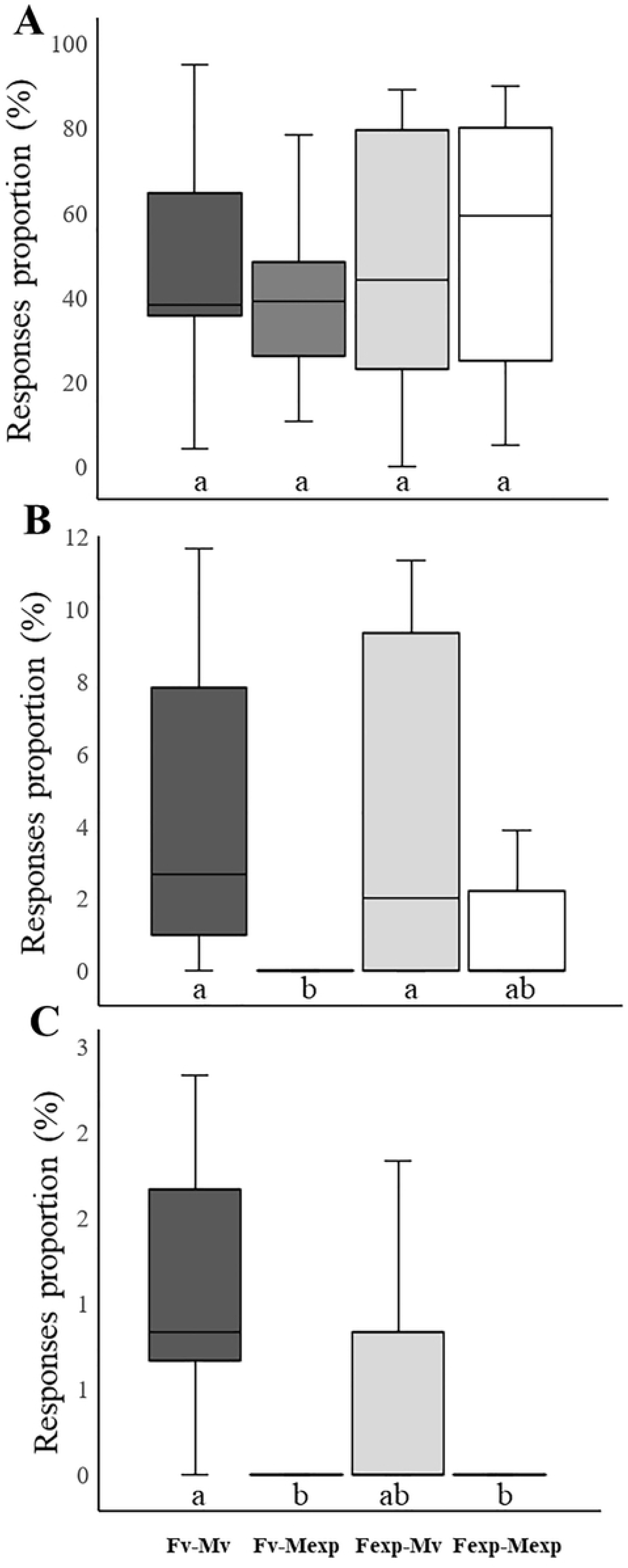
Behavioural mating responses of *Alphitobius diaperinus* in relation to gender and sexual experience. (A) Touching, (B) mounting and (C) copulation based on their gender and sexual experience. Different letters below the bars indicate significant differences according to the Mann-Whitney-U test with Bonferroni correction (α=0.05, p<0.05).

**Table 1.** Mating behaviour (in percentage; mean ± SD) of *Alphitobius diaperinus* adults based on their gender and sexual experience.

| Treatment | N | Touching | Mounting* | Copulation* |
| --- | --- | --- | --- | --- |
| Fv - Mv | 15 | 48.48 ± 24.77 | 5.75 ± 7.65 | 0.93 ± 0.73 |
| Fv - Mexp | 20 | 41.79 ± 23.01 | 0.08 ± 0.34 | 0.03 ± 0.11 |
| Fexp - Mv | 15 | 47.43 ± 28.73 | 4.28 ± 4.39 | 0.50 ± 0.85 |
| Fexp - Mexp | 15 | 49.81 ± 30.66 | 2.54 ± 6.69 | 0.13 ± 0.28 |

\*Significant difference (Kruskal-Wallis test, p<0.0001)

In the case of mating attempts, Fv-Mv and Fexp-Mv resulted in significantly longer attempts compared to Fv-Mexp (Fig 4a; Mann-Whitney-U test with Bonferroni correction, p<0.05); whereas we did not find differences in mating attempts in Fexp-Mexp compared to any other treatment. From the total mating attempts, no significant differences in unsuccessful and successful mating attempts among all the four treatments were found (Fig 4b and 4c; Kruskal-Wallis test, p>0.05). However, Mv was slightly more successful in mating with Fv compared to Fexp (Fig 4c). Moreover, from the total mating attempts showed by Fv-Mexp and Fexp-Mexp, only half was successful (Fig 4b and 4c, S1 Table).

**Fig 4.**
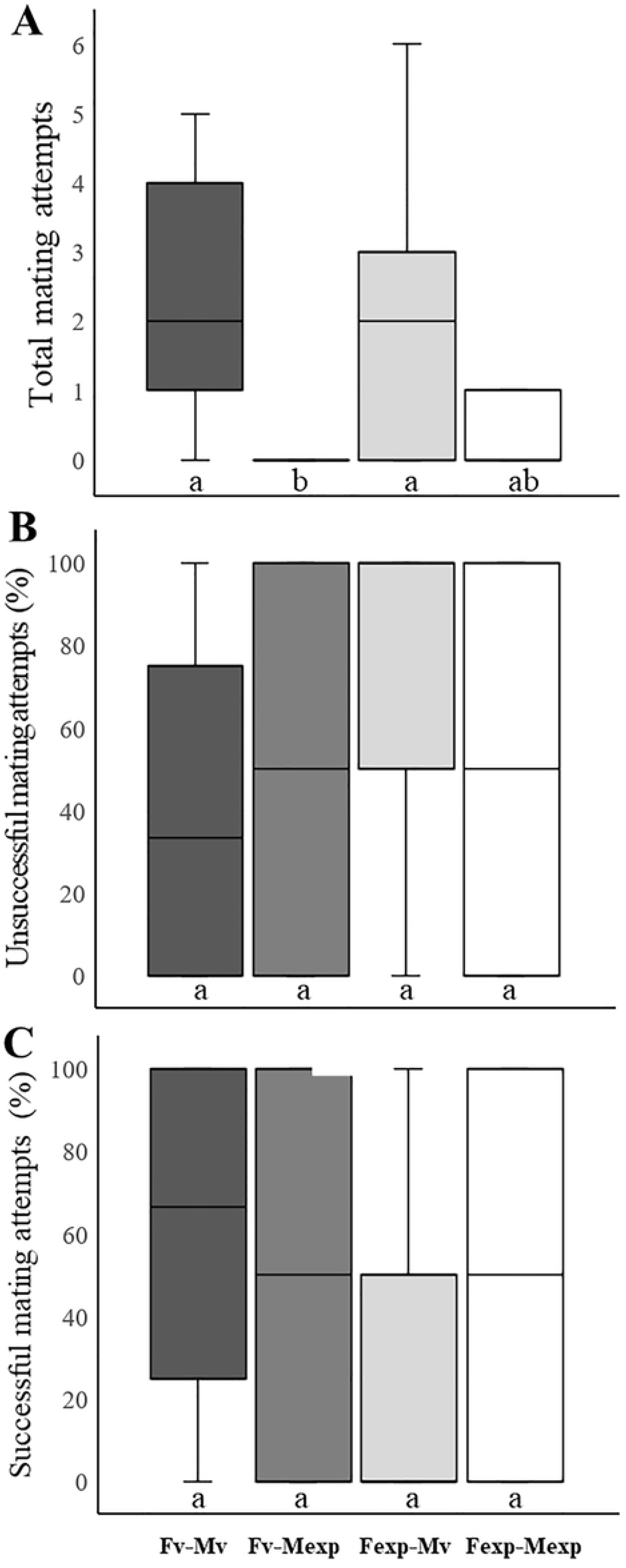
Mating attempts recorded of *Alphitobius diaperinus* adults based on their gender and sexual experience. (A) Total, (B) unsuccessful, and (C) successful mating attempts. Different letters below the bars indicate significant differences according to the Mann-Whitney-U test with Bonferroni correction (α=0.05, p<0.05).

### CHC profile of *A. diaperinus*

A total of 19 CHC compounds were obtained by GC-MS followed by a data curation (S2 Fig), which belonged to the following chemical groups alkanes, quinones, benzothiazoles, benzophenones, fatty acids and terpenes (Table 2). All the 19 identified compounds were detected in experienced adults but different concentrations, while only 14 compounds were detected in virgin adults. Among these CHC compounds, 2-methyl-1,4-benzoquinone and 2-ethyl-1,4-benzoquinone were detectable as significant components in the CHC profile for both males and females, although they were predominant in experienced adults (Table 2; S3 Fig). Interestingly, limonene, oleic acid, linoleic acid and nonacosane were identified only in experienced adults.

**Table 2.**
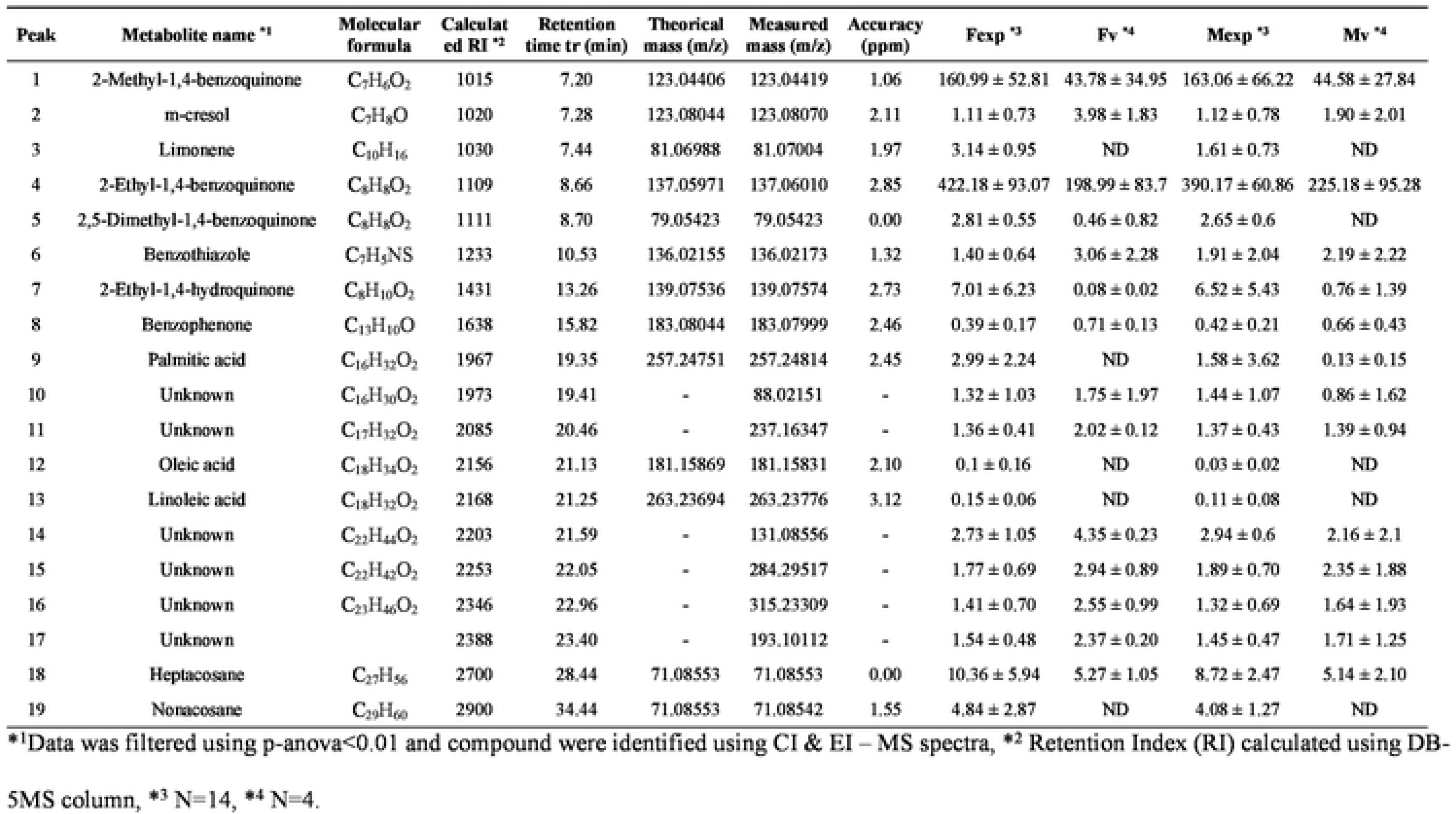
Mean quantity (ng/insect ± SD) of CHC compounds from adults of *Alphitobius diaperinus* based on gender and sexual experience.

| Peak | Metabolite name <sup>*1</sup> | Molecular formula | Calculated RI <sup>*2</sup> | Retention time tr (min) | Theoretical mass (m/z) | Measured mass (m/z) | Accuracy (ppm) | Fexp <sup>*3</sup> | Fv <sup>*4</sup> | Mexp <sup>*3</sup> | Mv <sup>*4</sup> |
| --- | --- | --- | --- | --- | --- | --- | --- | --- | --- | --- | --- |
| 1 | 2-Methyl-1,4-benzoquinone | C<sub>7</sub>H<sub>6</sub>O<sub>2</sub> | 1015 | 7.20 | 123.04406 | 123.04419 | 1.06 | 160.99 ± 52.81 | 43.78 ± 34.95 | 163.06 ± 66.22 | 44.58 ± 27.84 |
| 2 | m-cresol | C<sub>7</sub>H<sub>8</sub>O | 1020 | 7.28 | 123.08044 | 123.08070 | 2.11 | 1.11 ± 0.73 | 3.98 ± 1.83 | 1.12 ± 0.78 | 1.90 ± 2.01 |
| 3 | Limonene | C<sub>10</sub>H<sub>16</sub> | 1030 | 7.44 | 81.06988 | 81.07004 | 1.97 | 3.14 ± 0.95 | ND | 1.61 ± 0.73 | ND |
| 4 | 2-Ethyl-1,4-benzoquinone | C<sub>8</sub>H<sub>8</sub>O<sub>2</sub> | 1109 | 8.66 | 137.05971 | 137.06010 | 2.85 | 422.18 ± 93.07 | 198.99 ± 83.7 | 390.17 ± 60.86 | 225.18 ± 95.28 |
| 5 | 2,5-Dimethyl-1,4-benzoquinone | C<sub>8</sub>H<sub>8</sub>O<sub>2</sub> | 1111 | 8.70 | 79.05423 | 79.05423 | 0.00 | 2.81 ± 0.55 | 0.46 ± 0.82 | 2.65 ± 0.6 | ND |
| 6 | Benzothiazole | C<sub>7</sub>H<sub>5</sub>NS | 1233 | 10.53 | 136.02155 | 136.02173 | 1.32 | 1.40 ± 0.64 | 3.06 ± 2.28 | 1.91 ± 2.04 | 2.19 ± 2.22 |
| 7 | 2-Ethyl-1,4-hydroquinone | C<sub>8</sub>H<sub>10</sub>O<sub>2</sub> | 1431 | 13.26 | 139.07536 | 139.07574 | 2.73 | 7.01 ± 6.23 | 0.08 ± 0.02 | 6.52 ± 5.43 | 0.76 ± 1.39 |
| 8 | Benzophenone | C<sub>13</sub>H<sub>10</sub>O | 1638 | 15.82 | 183.08044 | 183.07999 | 2.46 | 0.39 ± 0.17 | 0.71 ± 0.13 | 0.42 ± 0.21 | 0.66 ± 0.43 |
| 9 | Palmitic acid | C<sub>16</sub>H<sub>32</sub>O<sub>2</sub> | 1967 | 19.35 | 257.24751 | 257.24814 | 2.45 | 2.99 ± 2.24 | ND | 1.58 ± 3.62 | 0.13 ± 0.15 |
| 10 | Unknown | C<sub>16</sub>H<sub>30</sub>O<sub>2</sub> | 1973 | 19.41 | - | 88.02151 | - | 1.32 ± 1.03 | 1.75 ± 1.97 | 1.44 ± 1.07 | 0.86 ± 1.62 |
| 11 | Unknown | C<sub>17</sub>H<sub>32</sub>O<sub>2</sub> | 2085 | 20.46 | - | 237.16347 | - | 1.36 ± 0.41 | 2.02 ± 0.12 | 1.37 ± 0.43 | 1.39 ± 0.94 |
| 12 | Oleic acid | C<sub>18</sub>H<sub>34</sub>O<sub>2</sub> | 2156 | 21.13 | 181.15869 | 181.15831 | 2.10 | 0.1 ± 0.16 | ND | 0.03 ± 0.02 | ND |
| 13 | Linoleic acid | C<sub>18</sub>H<sub>32</sub>O<sub>2</sub> | 2168 | 21.25 | 263.23694 | 263.23776 | 3.12 | 0.15 ± 0.06 | ND | 0.11 ± 0.08 | ND |
| 14 | Unknown | C<sub>22</sub>H<sub>44</sub>O<sub>2</sub> | 2203 | 21.59 | - | 131.08556 | - | 2.73 ± 1.05 | 4.35 ± 0.23 | 2.94 ± 0.6 | 2.16 ± 2.1 |
| 15 | Unknown | C<sub>22</sub>H<sub>42</sub>O<sub>2</sub> | 2253 | 22.05 | - | 284.29517 | - | 1.77 ± 0.69 | 2.94 ± 0.89 | 1.89 ± 0.70 | 2.35 ± 1.88 |
| 16 | Unknown | C<sub>23</sub>H<sub>46</sub>O<sub>2</sub> | 2346 | 22.96 | - | 315.23309 | - | 1.41 ± 0.70 | 2.55 ± 0.99 | 1.32 ± 0.69 | 1.64 ± 1.93 |
| 17 | Unknown |  | 2388 | 23.40 | - | 193.10112 | - | 1.54 ± 0.48 | 2.37 ± 0.20 | 1.45 ± 0.47 | 1.71 ± 1.25 |
| 18 | Heptacosane | C<sub>27</sub>H<sub>56</sub> | 2700 | 28.44 | 71.08553 | 71.08553 | 0.00 | 10.36 ± 5.94 | 5.27 ± 1.05 | 8.72 ± 2.47 | 5.14 ± 2.10 |
| 19 | Nonacosane | C<sub>29</sub>H<sub>60</sub> | 2900 | 34.44 | 71.08553 | 71.08542 | 1.55 | 4.84 ± 2.87 | ND | 4.08 ± 1.27 | ND |

<sup>\*1</sup>Data was filtered using p-anova<0.01 and compound were identified using CI & EI – MS spectra, <sup>\*2</sup> Retention Index (RI) calculated using DB-5MS column, <sup>\*3</sup> N=14, <sup>\*4</sup> N=4.

The (dis)similarity of CHC profile between the sexual experience groups of males and females of *A. diaperinus* was visually displayed by our NMDS analysis (Fig 5), which were highly congruent with a PCA also based on CHC profile (S2 Fig). Indeed, our PERMANOVA analyses did not show differences in CHC profiles based on gender identity (i.e., males vs females; Pseudo-F_(1, 35)_ = 0.757, p = 0.459), or within sexual experience groups (i.e., between virgins, between experienced adults; both p = 0.687). However, we did find statistical differences in CHC profiles according to the nested factor sexual experience (Pseudo-F_(2, 35)_ = 14.399, p<0.001). Specifically, we found that the Fexp CHC profile is different from that of Fv (PERMANOVA test with Holm correction, p<0.01) and Mv (PERMANOVA test with Holm correction, p<0.01). In contrast, Mexp were chemically different from Fv (PERMANOVA test with Holm correction, p<0.01) as well as from Mv (PERMANOVA test with Holm correction, p<0.01).

**Fig 5.**
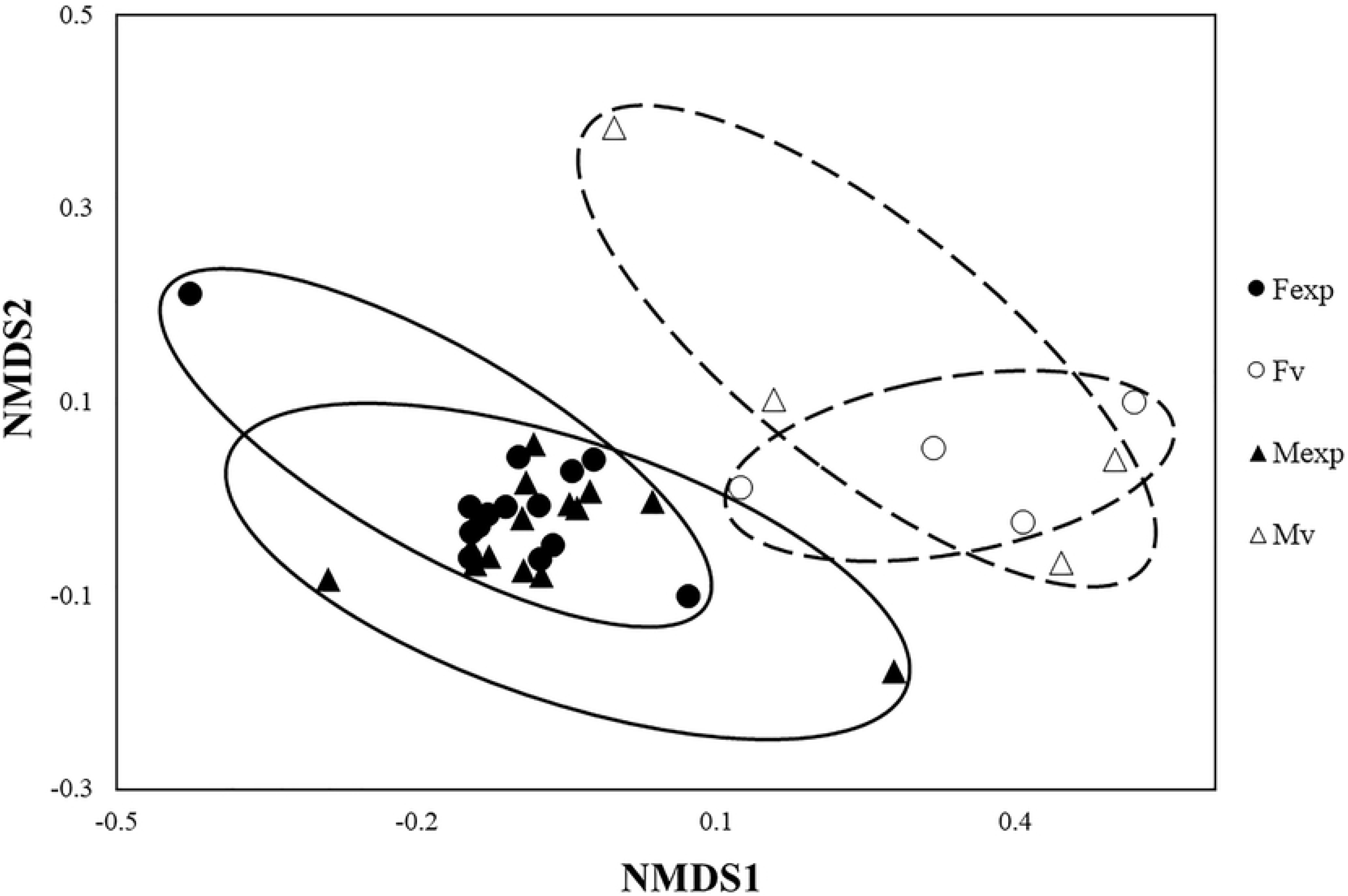
Comparison of *Alphitobius diaperinus* CHC profiles based on gender and sexual experience. A non-metric multidimensional scaling (NMDS) ordination based on Bray-Curtis similarities (stress=0.08036). Ellipses were drawn by hand.

## Discussion

The circadian cycles play an essential role in describing the mating behaviour of insect life (52–54), which might be considered a key to developing effective strategies for pest control (55,57). *Alphitobius diaperinus* seems not to show significant differences in mating activity along the day. However, a lower mating behaviour activity was observed at the end of the photophase and beginning of scotophase 13 h, which might be related to their cryptic behaviour when hiding from a threat or external factors (i.e., environmental changes) or to particular reproductive strategies of females to reduce her attractiveness and rejection to mate (5,56,57,68). In insects, the attractiveness of intra-specific odour relies on the interaction context (e.g., protection from environmental conditions, predators, food source or mate recognition), as recorded in some Curculionidae, Dermestidae, Tenebrionidae and other non-beetle species (37,55,59,68,80,91). In mating behaviour, the mate preference and attraction of males toward the odour of females can be based on their physiological changes and acquired experience (i.e., reproductive status and learned signal; (91)). For instance, in some Tenebrionidae species, experienced adults have contrasting mating behavioural responses compared to virgin adults (91,92). Interestingly, males and females of *A. diaperinus* were more attracted to the CHC profiles of both males and females although heavily dependent on the sexual experience condition, which can be related to the gregarious behaviour of this species and similarity of CHC profiles. Effectively, it has been shown that insects with similar odour profiles are less attractive to their gender counterparts compared to others with more different profiles (e.g., *Gryllodes sigillatus* (93)). Although it was previously pointed out that only virgin males released attractive compounds to attract virgin insects (e.g., aggregation pheromones; (5)), we have shown that CHC profiles of both males and females can be attractive to males and females, and their attractiveness relies on gender and sexual experience. Furthermore, we have shown that the insect cuticle diffusely releases attractive odour signals, in addition to the aggregation pheromones produced by exocrine glands in other beetles (94,95). In beetles, it is known the aggregation pheromones may be made by any sex (59,68,80,96). Unlike sex pheromones that act on only one sex, aggregation pheromones induce group formation of both sexes (37,43,55,68). In this regard, it may be evolutionarily advantageous for males to call females to reduce the time spent searching for a dispersed potential mate, and females have more chances to mate (80). Thus, it is clear that different combinations of odour profiles are necessary to attract males and females of *A. diaperinus* successfully. Thus, a single odour blend might not be enough.

Beetles during the premating phase show an antennal behaviour (59,65,80,97) related to the close-range assessment of potential mates, where male- or female produced contact pheromone is required for mating to be successful (58,61,62,67,80,98). Based on our observations, we speculate that both females or male adults of *A. diaperinus* release CHC compounds that allow the partner to be recognized by the opposite gender during antennation. This behaviour to recognise mates at short distances is mediated by semiochemicals such as a contact sex pheromone (58,61,62,67,98). In the copulatory phase, male beetles and *A. diaperinus* continue their antennation, touch with their maxillary palps and grasp the female’s cuticles with his prothoracic and mesothoracic leg (59,65,80,97) while mounting. On the leg, males possess tibial spurs and claws, which for other species facilitate grasping and control of females, preventing aggregated males from dislodging copulating males and distributing their own CHC profile overall the female’s cuticle (80). Thus, mate location and sex discrimination, the first steps of mating, is maybe due to a combination of physical and chemical recognition by the antenna and legs (59).

It is usually expected that adult males tend to mate with any available female. However, we have often recorded low mating activity of Mexp with females, compared to Mv, which could be explained by the sexual conflict (99,100). To increase reproductive success, females mate with multiple males to select their best candidate (99), giving the later males more chances to succeed. Experienced males have already mated and might be more selective to mate; therefore, they would prefer to mate with Fexp, which are less likely to have new mating partners than Fv. Moreover, the sexual experience would also reduce the amount of time in Mexp mounting and copulation since they are shorter with Fexp than Fv. Unlike Mexp, Mv has a natural-sexual need to copulate with a mate. Therefore, Fv and Fexp are both attractive to Mv, although Fexp seems to be more attractive. This behaviour is also known in other insects, where virgin males mate indiscriminately with females and do not recognise their sexual experience (101,102).

The CHC play an essential role in chemical communication during the mating behaviour of insects (49,58,60–62,67,70,72–74). According to our results, differences in CHC profiles of *A. diaperinus* adults are mainly based on their sexual experience and not gender, which explains why the odour profile of both genders can be attractive. Moreover, CHC profiles can explain the results of our mating bioassays and suggest that CHC compounds are the key to mate choice and reproduction in *A. diaperinus*. CHC profile similarity between males and females can also explain the expected homosexual mating behaviour observed in *A. diaperinus* and diverse beetle species (77,103,104). Thus, the CHC profile of the sexual experience in adults serves as a crucial species-specific mating cue, but not for sex discrimination (72). Further studies are being carried out to identify the chemical basis of *A. diaperinus* attractiveness.

## Acknowledgments

All the authors thank to “The Max Planck Partner Group” for their financial support. ECQ doctoral studies were supported by a doctoral scholarship through the “Proyecto de Mejoramiento y Ampliación de los Servicios del Sistema Nacional de Ciencia Tecnología e Innovación Tecnológica (8682-PE)” funded by the World Bank, CONCYTEC, and FONDECYT (Doctoral Program 010-20180). CM was kindly supported by a research grant from “Fondo Nacional de Desarrollo Científico (CONVENIO N° 126-2020-FONDECYT)”.

## Supporting information

**S1 Fig. Circadian pattern of sexual activity of *Alphitobius diaperinus* based on their sexual condition (virgin pair and experienced pair).** Circadian pattern test: N=10 (= 100%) for each trial observed at one-hour intervals for a 24-h period during five days in the laboratory (H1–H12: 07:00–19:00 and H13–H24:19:00–07:00). Significant difference according to the Mann-Whitney-U test with Bonferroni correction (α=0.05, p<0.05).

**S1 Table. Mating behaviour (in percentage; mean ± SD) and mating attempts recorded of *Alphitobius diaperinus* based on their gender and sexual condition.**

**S2 Fig. Biplot of principal component analysis (PCA) of curated compound-data matrix shows the PCA scores and loadings plot from adults of *Alphitobius diaperinus*.** (A) PC1 vs PC2, (B) PC2 vs PC3. The peaks represent metabolites described in Table 2.

**S3 Fig. GC-MS (CI) chromatograms of adults of *Alphitobius diaperinus* based on their gender and sexual condition.** (A) Fv, (B) Fexp, (C) Mv, (D) Mexp. 1) MBQ, 2) m-cresol, 3) limonene, 4) EBQ, 5) 2,5-dimethyl-1,4-benzoquinone, 6) benzothiazole, 7) 2-ethyl-1,4-hydroquinone, 8) benzophenone, 9) palmitic acid, 10) unknown, 11) unknown, 12) oleic acid, 13) linoleic acid, 14) unknown, 15) unknown, 16) unknown, 17) unknown, 18) heptacosane, 19) nonacosane. IS, internal standard, C16.

